# First person experience of threat modulates cortical network encoding human peripersonal space

**DOI:** 10.1101/314971

**Authors:** A.W. de Borst, M.V. Sanchez-Vives, M. Slater, B. de Gelder

## Abstract

Peripersonal space is the area directly surrounding the body, which supports object manipulation and social interaction, but is also critical for threat detection. In the monkey, ventral premotor and intraparietal cortex support initiation of defensive behavior. However, the brain network that underlies threat detection in human peripersonal space still awaits investigation. We combined fMRI measurements with a preceding virtual reality training from either first or third person perspective to manipulate whether approaching human threat was perceived as directed to oneself or another. We found that first person perspective increased body ownership and identification with the virtual victim. When threat was perceived as directed towards oneself, synchronization of brain activity in the human peripersonal brain network was enhanced and connectivity increased from premotor and intraparietal cortex towards superior parietal lobe. When this threat was nearby, synchronization also occurred in emotion-processing regions. Priming with third person perspective reduced synchronization of brain activity in the peripersonal space network and increased top-down modulation of visual areas. In conclusion, our results showed that after first person perspective training peripersonal space is remapped to the virtual victim, thereby causing the fronto-parietal network to predict intrusive actions towards the body and emotion-processing regions to signal nearby threat.

## Introduction

Humans initiate many protective behaviors in daily life (e.g. avoiding sharp objects, collision), as well as in more extreme situations (e.g. avoiding physical aggression). The fact that the body is the prime tool for self-protection attributes a special status to the representation of the space directly surrounding the body within reach of the limbs, the so-called peripersonal space (PPS). The brain has a specialized system for processing information within the PPS, consisting of two interconnected multisensory regions, the ventral premotor cortex (vPM) and the ventral intraparietal area (VIP) (Graziano and Cooke, 2006). Visuo-tactile neurons in the vPM and VIP of the monkey brain are specifically activated by stimuli within the PPS, with matched visual and tactile receptive fields (Rizzolatti et al., 1981a; Rizzolatti et al., 1981b; Colby et al., 1993; Graziano and Gross, 1993; Graziano et al., 1994; Graziano et al., 1997; Duhamel et al., 1998; Graziano, 1999). The alignment between the visual and tactile receptive fields allows for the encoding of multisensory space in a common reference frame (Fogassi et al., 1992; Graziano et al., 1997). Similar to the neurophysiological work in monkeys, research indicates that the human PPS encoding network consists of multisensory regions in vPM, intraparietal sulcus (IPS) and, possibly, primary somatosensory cortex (PSC) (Sereno and Huang, 2006; Makin et al., 2007; Gentile et al., 2011; Brozzoli et al., 2012a; Brozzoli et al., 2012b; Huang et al., 2012; Brozzoli et al., 2013; Holt et al., 2014; Ferri et al., 2015).

Research in monkeys has found that these multisensory regions representing peripersonal space support object manipulation (Maravita et al., 2003; Bremmer, 2005). More recently it has been shown that electrical stimulation of the neurons in the macaque vPM and VIP causes the monkey to display defensive behavior similar to sustained air puff evoked responses (Cooke and Graziano, 2003, 2004). These results suggest that the peripersonal space also functions as an area of safety around the body and that the underlying brain regions allow for the initiation of defensive behavior towards threats (Cooke and Graziano, 2003; Graziano and Cooke, 2006). In humans, behavioral work has further supported the hypothesis that peripersonal space performs an important defensive function.

For example, peripersonal space is enlarged in people with claustrofobic fear and in schizophrenic patients (Lourenco et al., 2011; Holt et al., 2015). Other work has shown a relationship between threatening objects and nearby space. For example, reachable space is reduced when threatening objects are pointed towards the participants (Coello et al., 2012). De Haan et al. (2016) have shown that people with fear for spiders have shorter tactile reaction times to near-by approaching spiders than butterflies. However, it is still unknown whether in humans these effects are due to changes in the peripersonal brain network, or whether other brain regions underlie these effects.

In this study we researched the defensive function of peripersonal space and its underlying brain network in humans. In order to create a relevant and realistic stimulus we used immersive virtual reality. Immersive virtual reality typically creates the perceptual illusion of ‘presence’, the illusion of being in the displayed scene (Sanchez-Vives and Slater, 2005), and may elicit ‘plausibility’, which is the illusion that the events in the scenario are really occurring (Slater, 2009). These illusions, together with body ownership - the illusion of owning a body (part) other than one’s own (Blanke, 2012) - and first person perspective, typically lead participants to behave similarly in VR to how they would behave in a comparable situation in reality (Yee et al., 2009; Banakou et al., 2013; Maister et al., 2013; Peck et al., 2013; Maister et al., 2015; Banakou et al., 2016). The human brain network underlying body ownership is strongly linked to the peripersonal space brain network, sharing several brain regions (Grivaz et al., 2017). Perspective may also have an important function in social aspects of peripersonal space. Brozzoli et al. (2013) suggested the presence of a shared PPS representation between the self and others in ventral PM, which could support social interactions. It is unclear however, whether the observed PPS effects in PM arise because the first person perspective of the other’s hand leads to displacement of the own PPS to the other’s (in line with Oosterhof et al., 2012), or because the representation of space surrounding the other’s hand activates the PPS network (in line with Schaefer et al., 2012).

Here, we investigated how priming with whole-body first person or third person perspective virtual reality training modulated the PPS network during subsequent 3D perception of approaching social threat. The fMRI design of this study follows earlier investigations using free viewing of natural scenes (Bartels and Zeki, 2004; Hasson et al., 2004), rather than repeated presentation of static stimulus conditions. During the perception of natural scenes, when many aspects of the scene have to be processed simultaneously, functional specialization remains intact (Bartels and Zeki, 2004) and individual brains show highly synchronous activity (Hasson et al., 2004). In order to control for all lower level stimulus properties between conditions, we presented an identical 3D video during fMRI measurements in both conditions, while participants were primed with a first or third person perspective using the preceding virtual reality training. We hypothesize that activation in PM, IPS and PSC will be stronger when the observer is primed to inhabit the space of the virtual victim (first person perspective) than when the observer is primed to passively perceive the threat in the virtual victim’s space (third person perspective). Furthermore, on the basis of monkey research, we expect visual and somatosensory regions to send information to PM and IPS in order to initiate defensive behavior. We hypothesize that the supramarginal gyrus/temporoparietal junction (SMG/TPJ) will be involved during peripersonal space intrusion, given that it was found to be part of the PPS network (Grivaz et al., 2017), but its exact role will need to be investigated. We expect stronger activation of emotion-related structures, such as amygdala (AMG), during first person perspective experience of nearby threat, as spatial proximity is an important factor for fear-induced defensive responses (Ahs et al., 2015).

## Materials and Methods

### Participants

Twenty healthy volunteers of Dutch and German nationality participated in this study. Half of the participants were male (mean age 22.3 years; range 19-28) and half were female (mean age 20.3 years; range 18-24). All participants had normal or corrected-to-normal vision and gave their informed consent. All participants were fluent in Dutch. Exclusion criteria were the institute’s MRI safety criteria. Due to the nature of the stimuli, we also excluded volunteers who had a criminal record, or a history of physical or emotional abuse. The study was approved by the local ethical committee.

### Stimuli and materials

In this study we used virtual reality training and 3D videos. Immersive virtual reality typically creates the perceptual illusion of ‘presence’, the illusion of being in the displayed scene (Sanchez-Vives and Slater, 2005), and may elicit ‘plausibility’, which is the illusion that the events in the scenario are really occurring (Slater, 2009). These illusions, together with body ownership and first person perspective, typically lead participants to behave similarly in VR to how they would behave in a comparable situation in reality (Yee et al., 2009; Banakou et al., 2013; Maister et al., 2013; Peck et al., 2013; Maister et al., 2015; Banakou et al., 2016). The stimuli consisted of a VR scenario, two auditory stimuli containing instructions, and one 3D split-screen video. The VR scenario displayed a female avatar in the hallway of a house (see Fig. 1, left). There were two mirrors in the hallway, two doors on opposite ends, and a sideboard. The scenario could be viewed from a first person perspective (1PP), or a third person perspective (3PP). The VR environment was built in Unity (Unity Technologies, San Francisco, USA). The participants viewed the VR scenario using an Oculus Rift DK2 (Oculus VR, Menlo Park, USA), which is a head-mounted display especially designed to view VR. The Oculus Rift has an OLED display with a 960 x 1080 resolution per eye, and uses an infrared camera for positional tracking of the headset. Stereoscopic vision was obtained by projecting the stimulus in a slightly different angle to the left and right eye. Each of the auditory stimuli lasted 2:08 min and consisted of a female voice giving instructions for several visuomotor exercises in the VR scenario from either the first person or the third person perspective (e.g. “move your head to the right until you see the edge of the mirror”). The 3D split-screen video shown in the MRI scanner was a recording of a VR domestic abuse scenario from first person perspective (Fig. 1, right). In this abuse scenario, a male avatar entered the hallway from one of the doors and started addressing the female avatar in a demeaning and aggressive manner. Over the course of 2:37 min, the male avatar throws the phone through the hallway and approaches the female closely while continuing to verbally abuse her (for a transcription of the narrative see Supplementary Materials, Table S2). The 3D video was viewed inside the MRI scanner using VisStim MRI-compatible goggles (Resonance Technology, Northridge, USA). The VisStim goggles contain two displays, each with a 600 x 800 resolution, set within a rubber head mount. Similar to the Oculus Rift, stereoscopic vision was obtained by projecting the split-screen video onto the two screens. A VR questionnaire consisting of 17 items relating to different aspects of the virtual reality VR experience was used as well.

**Fig. 1.**
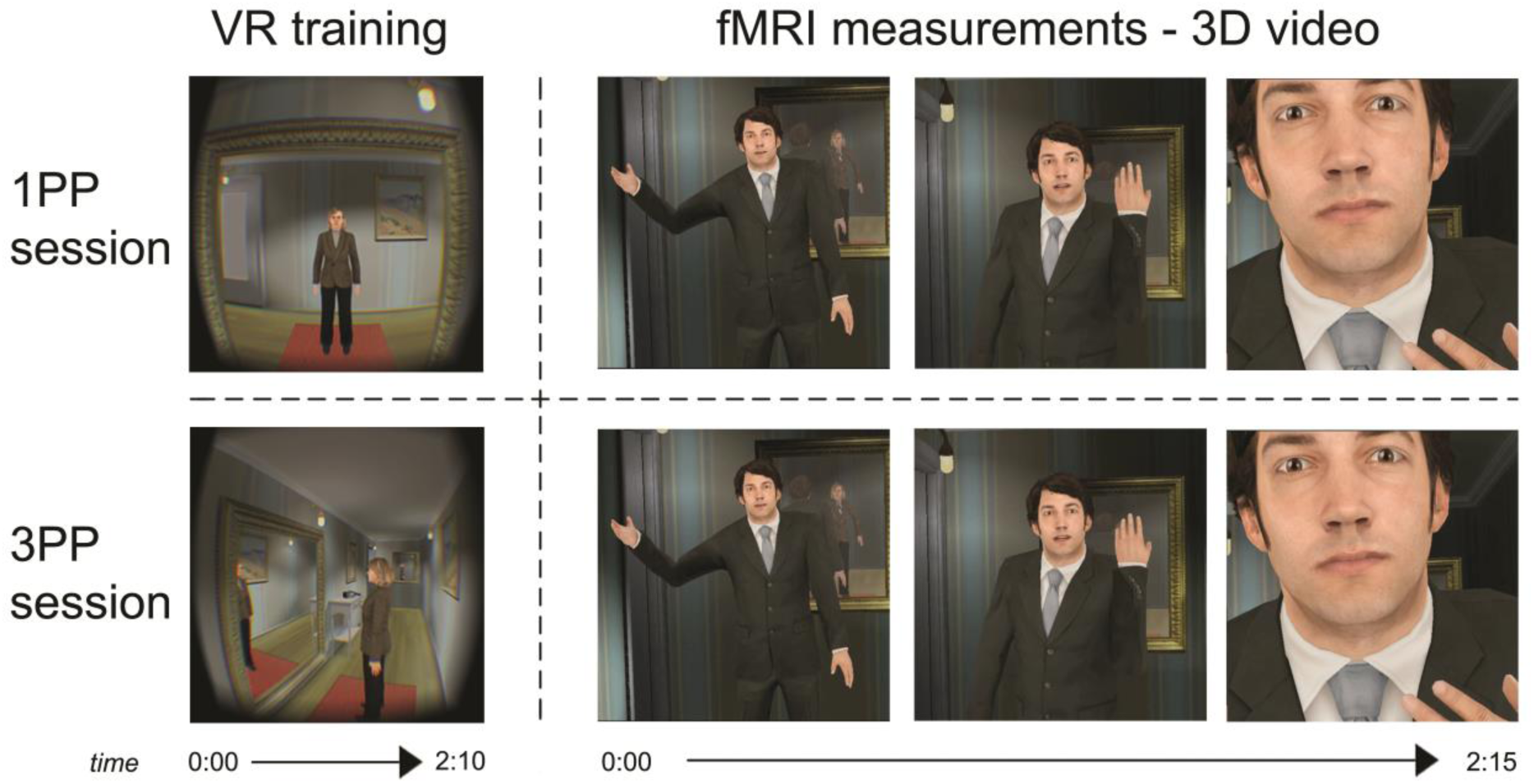
Experimental design. In two counterbalanced sessions, participants were immersed in a virtual reality scenario (VR training) where they either observed the scenario from a first person perspective and performed visuomotor exercises congruent with the female character’s movements (1PP; top left), or observed the scenario from a third person perspective and performed visuomotor exercises congruent with the virtual camera’s movements (3PP; bottom left). In both sessions, participants were subsequently moved to an MRI scanner, where they watched a continuous 3D video showing a social threat situation in which a male aggressor approached the viewer and entered their peripersonal space (right).

### Experimental procedure

The experiment consisted of two different sessions, which were one week apart. In the first session, participants were informed about the study and signed the informed consent form. Afterwards, the subjects were familiarized with the MRI environment. Subsequently, outside of the scanner room, they put on the Oculus Rift and followed auditory instructions to perform several visuomotor exercises. They saw the VR scenario from either the first-person perspective or the third-person perspective (counterbalanced). In the first-person perspective (1PP session) they looked into a full-length mirror and saw the female avatar performing head movements that were consistent with the participants’ own movements, thereby contributing to the illusion of body ownership (Banakou and Slater, 2014) (Fig. 1, top left). In the third-person perspective scenario (3PP session), the participants performed the same movements as indicated by the auditory instructions, but instead they viewed the mirror and female avatar from a slight distance and did not have a virtual body (Fig. 1, bottom left). In this perspective, the virtual camera viewpoint, rather than the female avatar, moved consistently with the participants’ head movements. During the training the participants were not exposed to the virtual threat. After the first person or third person visuomotor training with the Oculus, the participants were blindfolded (in order to maintain the 1PP or 3PP illusion) and led to the MRI scanner. During fMRI measurements the participants passively viewed a 3D video of the VR environment where they, from the perspective of the female character, were verbally and psychologically abused by an approaching male character (Fig. 1, right). The 3D video shown during fMRI measurements was *identical* in both sessions. At the end of the first session, participants filled out the VR questionnaire. At the end of the first session participants were partially debriefed about the study, were asked about how they experienced the scenario and if they were affected by it. Moreover, they were asked to contact the experimenter if they had any reoccurring thoughts or feelings about the experiment.

After a week, participants came back to the lab and followed the same procedure as during the first session, but during this second session they viewed the VR scenario from the other perspective (e.g. if they viewed it from third person perspective in the first session, then they viewed it from the first person perspective in the second session). All other aspects of the session were identical. They also filled out the VR questionnaire again at the end of the session and were debriefed about the contents and meaning of the study. Again, the emotional state of the participants was assessed and they were asked to contact the experimenter if they had any reoccurring thoughts or feelings, or were otherwise affected by participating in this experiment. No participant reported to be distressed by the experiment or have persisting thoughts or feelings about the experiment.

### Design

The order of the VR training perspective (1PP vs. 3PP) between sessions was counterbalanced across subjects, so that half of the males and half of the females had the session order 1PP-3PP and the other halves had the opposite order. The 3D video that was watched during fMRI measurements was identical in both sessions and was preceded and followed by 3 seconds of fixation. The two experimental conditions were the perception of the 3D video preceded by 1PP VR training (1PP session) and the perception of the 3D video preceded by 3PP VR training (3PP session). The 3D stimulus was shown once in each session, similar to other naturalistic research (e.g. Hasson et al., 2004)

### Data acquisition

A 3T Siemens MR scanner (MAGNETOM Prisma, Siemens Medical Systems, Erlangen, Germany) was used for imaging. Functional scans were acquired with a multiband gradient echo echo-planar imaging sequence with a Repetition Time (TR) of 1500 milliseconds (ms) and an Echo Time (TE) of 30 ms. The functional run consisted of 90 volumes comprising 57 slices (matrix = 800×800, 2 mm isotropic voxels, inter slice time = 26 ms, flip angle = 77°). After the functional run, high resolution T1-weighted structural images of the whole brain were acquired with an MPRAGE with a TR of 2250 ms and a TE of 2.21 ms, 192 slices (matrix = 256×256, 1 mm isotropic voxels, flip angle = 9°).

### Statistical analyses

#### Questionnaire analyses

The VR questionnaire contained questions relating to the subjective experience of the 3D social threat video. The scores on the VR questionnaire were compared between sessions (Session; 1PP vs. 3PP) by conducting an ordinal logistic regression analysis, with Score as an independent variable and Session as factor, thresholded at p < 0.05 (Fig. 2). From the VR experience questionnaire, we used the scores on the question “To what extent have you experienced the situation as if it was real?”, the question “To what extent did you feel in the female body and lived the situation as if you were the woman?” and the question “To what extent did you feel identified with the female body during the experience?” to analyze, respectively, the perceived Plausibility, Body Ownership and Identification during the viewing of the 3D social threat video.

**Fig. 2.**
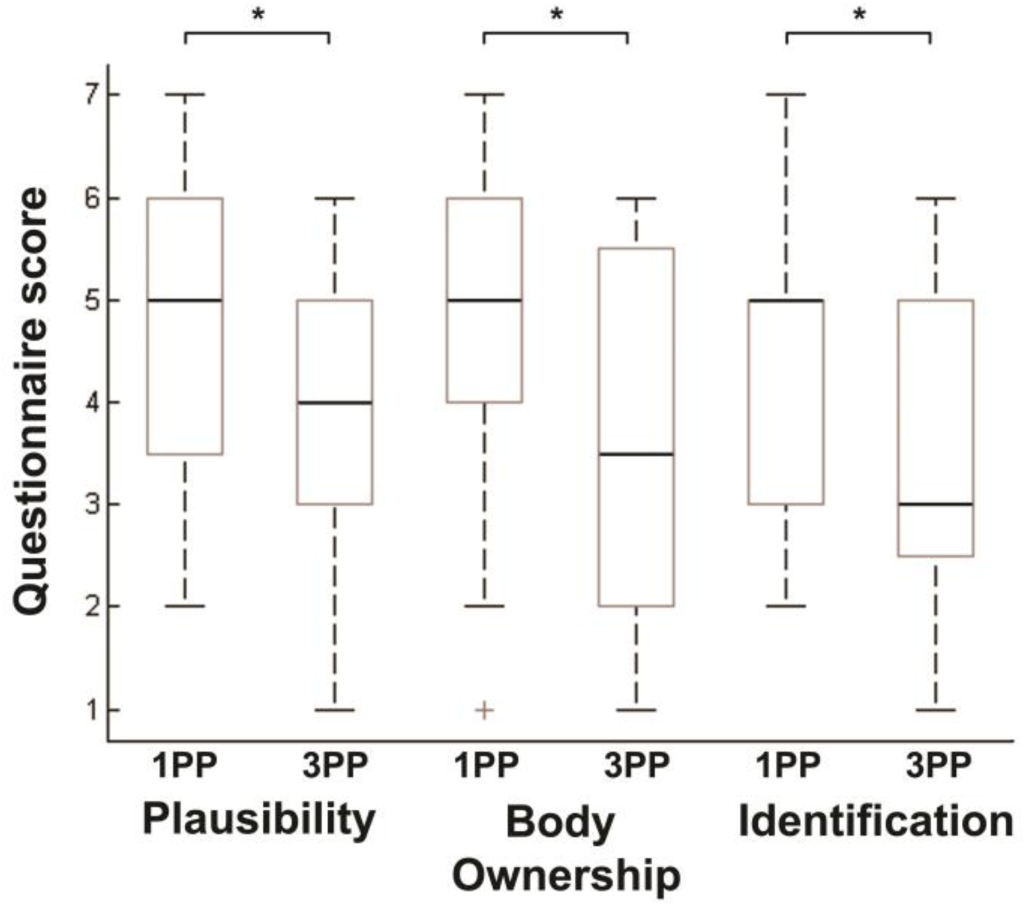
Boxplots of VR questionnaire data. The boxplots show the medians, interquartile ranges, maximum and minimum (as indicated by the stems) and outliers of the questionnaire scores that addressed the subjective experience of Plausibility, Body Ownership and Identification. For each question the results for the 1PP session are displayed on the left and those of the 3PP session on the right. An asterisk indicates a significant difference (p < 0.05).

#### Functional MRI pre-processing

The fMRI data were pre-processed and visualized using fMRI analysis and visualization software BrainVoyager QX version 2.8.4 (Brain Innovation B.V., Maastricht, the Netherlands). Functional data were corrected for head motion (3D motion correction, sinc interpolation), corrected for slice scan time differences and temporally filtered (high pass, GLM-Fourier, 2 sines/cosines). For the ISC analyses, it is recommended to spatially smooth the functional data with a Gaussian smoothing kernel of slightly larger than double the original voxel size (Pajula and Tohka, 2014). Therefore, the functional data was spatially smoothed using a Gaussian kernel with a FWHM of 5 mm. The anatomical data were corrected for intensity inhomogeneity (Goebel et al., 2006) and transformed into Talairach space (Talairach and Tournoux, 1988). The functional data were then aligned with the anatomical data and transformed into the same space to create 4D volume time-courses (VTCs).

#### Anatomical mask

For the ISC analyses, we used an anatomical mask, which included several probabilistic cytoarchitectonic regions-of-interest, covering most of the cortex (see Supplementary Materials, Figure S1 and Table S1). Statistical Parametric Mapping (SPM; version 12, Functional Imaging Laboratory, London, UK) was used to extract probabilistic cytoarchitectonic maps from the SPM Anatomy Toolbox (version 1.8, Forschungszentrum Jülich GmbH; Eickhoff et al., 2005). Each voxel in a probabilistic region reflects the cytoarchitectonic probability (10–100%) of belonging to that region. We followed a procedure to obtain maximum probability maps as described in Eickhoff et al. (2006), as these are thought to provide ROIs that best reflect the anatomical hypotheses. This meant that all voxels in the ROI that were assigned to a certain area were set to “1” and the rest of the voxels were set to “0”. The ROIs were transformed from MNI space to Talairach space (as Talairach was used in the other analyses). We extracted the Colin27 anatomical data to help verify the subsequent transformations. In order to transform the ROIs and the anatomical data from MNI space to Talairach space, we imported the ANALYZE files in BrainVoyager, flipped the x-axis to set the data to radiological format, and rotated the data −90° in the x-axis and +90° in the y-axis to get a sagittal orientation. Subsequently, we transformed the Colin27 anatomical data to Talairach space (Talairach and Tournoux, 1988; Goebel et al., 2006) and applied the same transformations to the cytoarchitectonic ROIs (see also de Borst and de Gelder, 2016). The bilateral probabilistic cytoarchitectonic maps of the AMG, insula (INS) and anterior cingulate cortex (ACC) were used as region-of-interest (ROI) in the ISC time course analysis (see section below and Figure 5).

**Fig. 3.**
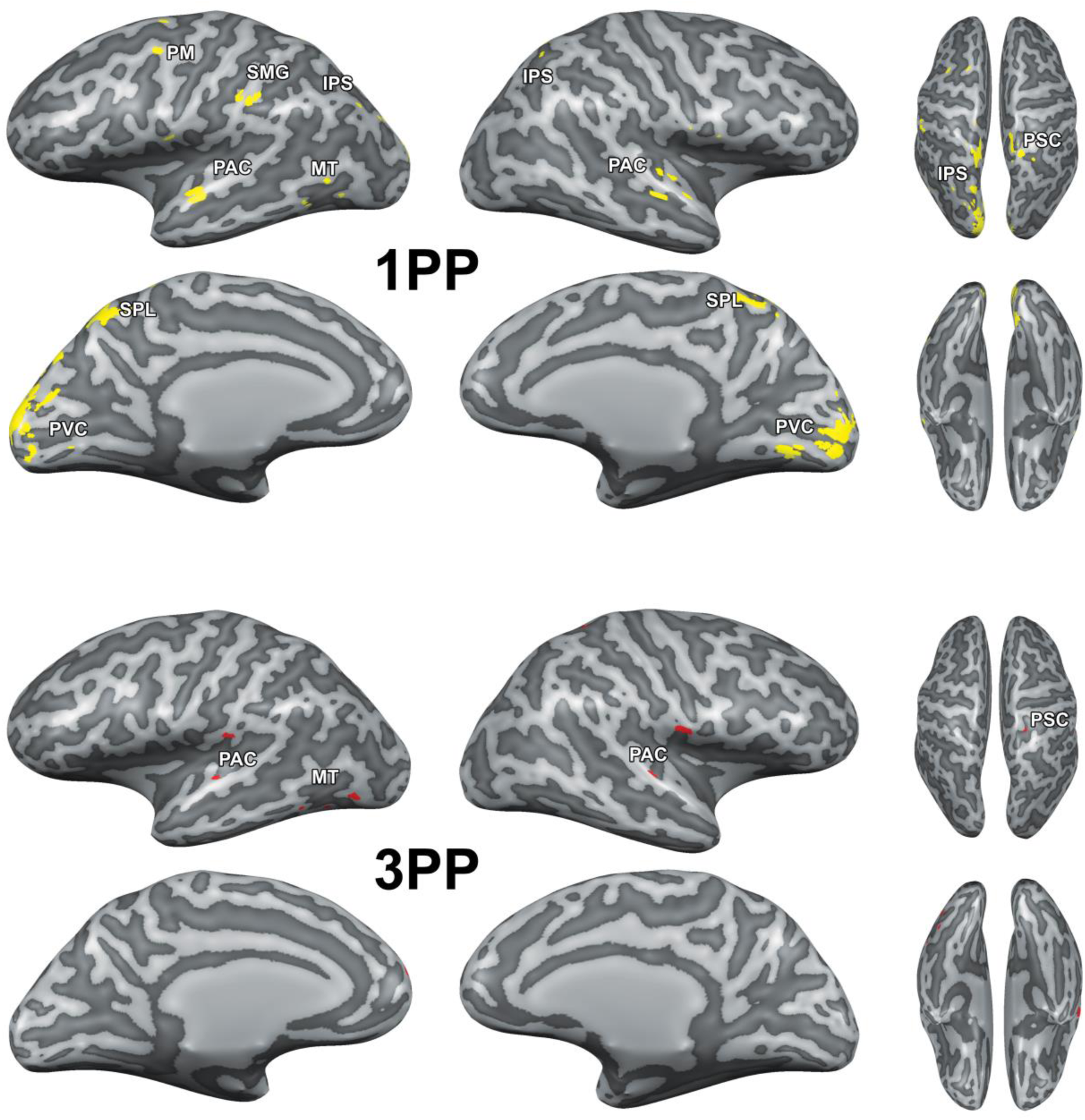
Intersubject correlation differences (permutation testing, N = 20, p[corrected] < 0.05) between first (1PP) and third person perspective (3PP) primed perception of an identical 3D threat video. Voxels that showed significantly higher ISC after 1PP training than 3PP training are indicated in yellow (top). Voxels that showed significantly higher ISC after 3PP training than 1PP training are indicated in red (bottom). PAC = primary auditory cortex, PM = premotor cortex, SMG = supramarginal gyrus, MT = middle temporal area, SPL = superior parietal lobe, PVC = primary visual cortex, IPS = intraparietal sulcus, PSC = primary somatosensory cortex.

**Fig. 4.**
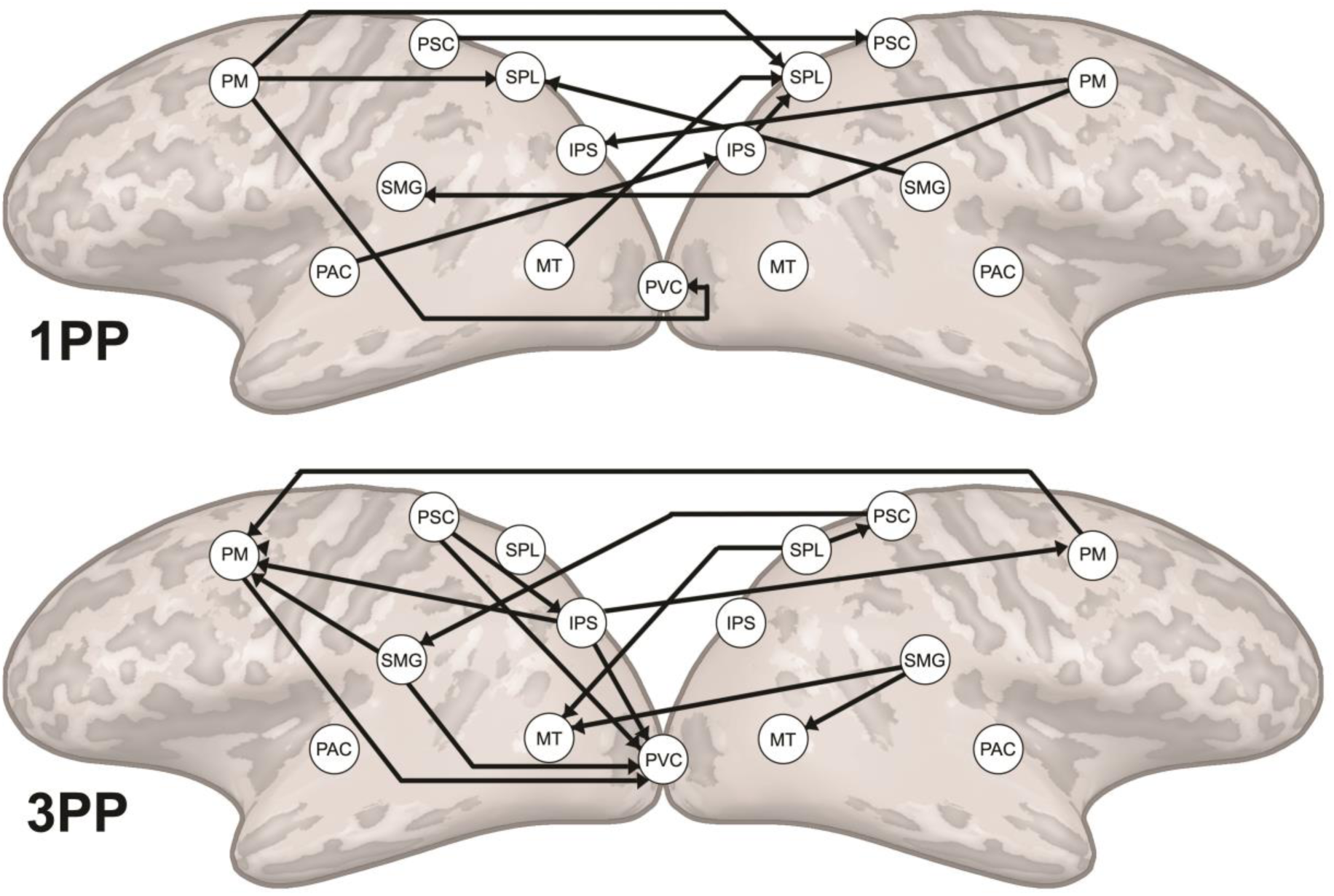
Differences in effective connectivity between first (1PP) and third person perspective (3PP) primed perception of an identical 3D threat video (RFX ANCOVA, N = 20, p[corrected] < 0.05). The arrows indicate the direction of the connectivity between regions that is unique for each condition. Abbreviations as in Fig. 3.

**Fig. 5.**
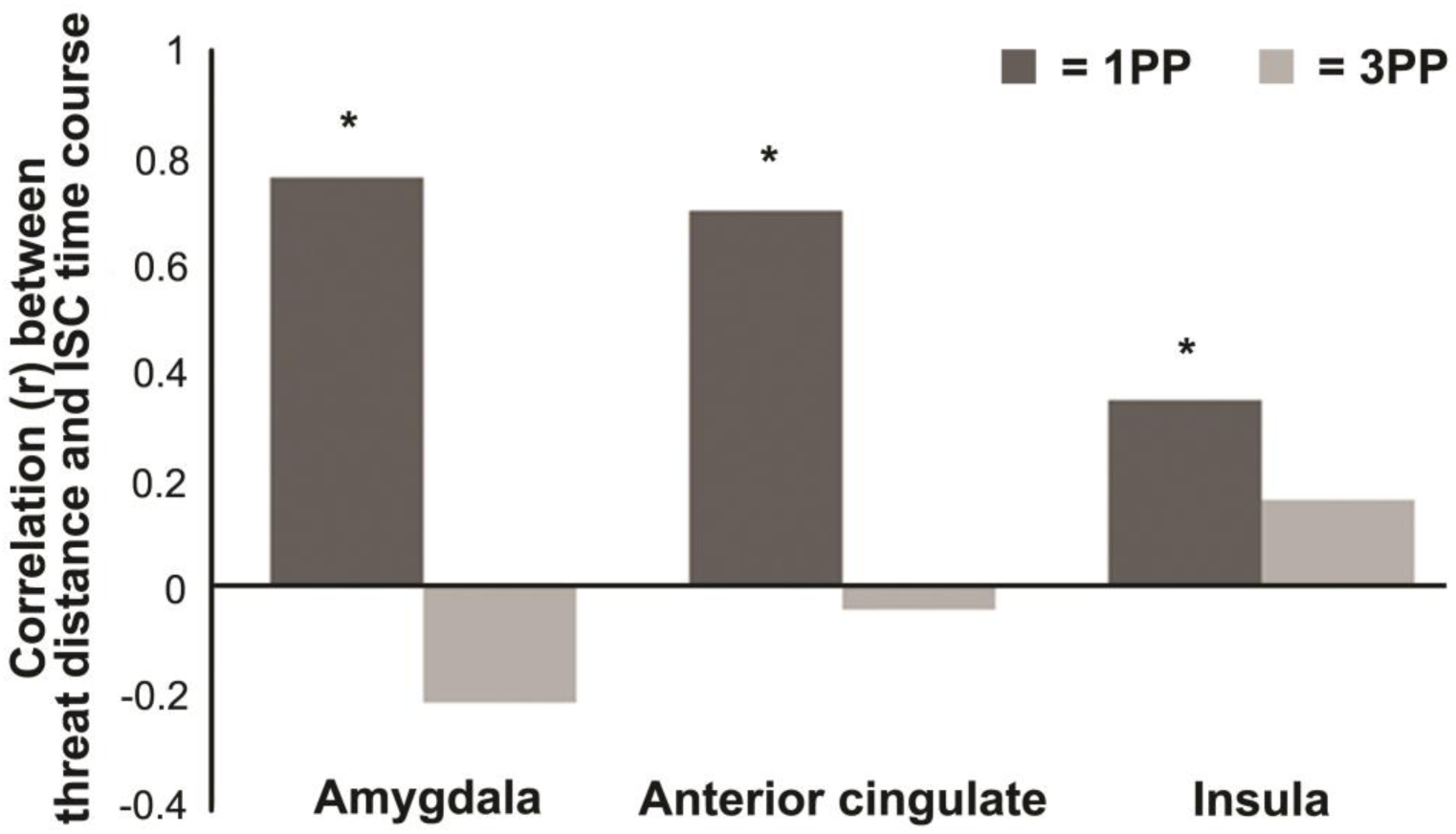
Correlation between threat distance and the intersubject correlation (ISC) time course within each region-of-interest in the 1PP (dark grey) and the 3PP (light grey) session. An asterisk indicates a significant relationship between threat distance and ISC.

#### Intersubject correlation

The ISC toolbox for fMRI in Matlab (Kauppi et al., 2014) and in-house Matlab scripts were used for ISC analyses with Matlab version R2013b, 8.2.0.701 (The MathWorks Inc., Natick, USA). We calculated ISC difference maps to reveal significant differences between conditions. Pearson’s correlation coefficient was used to calculate voxel-wise temporal correlations between all possible subject pairs (N (N - 1) / 2) for all points of the VTC time-course (90 volumes), or selected time windows (10 volumes). The ISC difference maps between the sessions were calculated as described in Kauppi et al. (section 2.2.4; 2014). In the ISC difference analyses a Fisher’s z-transformation (ZPF; Fisher, 1915) was applied to each pairwise correlation in each voxel. Subsequently, a sum ZPF statistic for the difference between the two conditions (1PP and 3PP) was calculated over all subject pairs of one group and tested against the null hypothesis that each ZPF value comes from a distribution with zero mean (no difference between 1PP and 3PP) using non-parametric permutation testing. The null distribution was obtained by randomly flipping the sign of pairwise ZPF statistics before calculating the sum ZPF statistic using 25000 permutations. Maximal and minimal statistics over the entire image corresponding to each labeling were saved. The map was thresholded at α = 0.05 using the permutation distribution of maximal statistic, which accounts for the multiple comparisons problem by controlling the FWER (see Nichols and Holmes, 2002).

In a second set of analyses we calculated the dynamic ISC of brain activity by computing the average ISC for each acquired volume using a 10-sample moving average in three ROIs (AMG, INS, ACC). This approach resulted in 81 ISC maps, each reflecting the moment-to-moment degree of intersubject synchronization across participants. The mean across all voxels of each ROI was calculated for each of the time points, resulting in a time course of ISC for each ROI. The first 5 volumes and the last 3 volumes were removed from each time course. The resulting time courses (consisting of 73 time points) were correlated with a box-car function with “0” for the first 34 time points that corresponded to the time during which the aggressor was in far space, and “1” for the remaining 39 time points that corresponded to the time during which the aggressor was in peripersonal space. The correlation coefficient was tested against the null hypothesis that there is no relationship between the observed phenomena and thresholded at p < 0.05.

#### Granger Causality Mapping

We used the RFX GCM plugin of BrainVoyager QX 2.8 and in-house Matlab scripts to calculate effective connectivity between brain regions. Granger Causality Mapping (GCM) (Roebroeck et al., 2005) uses vector autoregressive models of fMRI time series in the context of Granger causality. A time-series of voxel X Granger causes a time-series of voxel Y, if the past of X improves the prediction of the current value of Y, given that all other relevant sources of influence have been taken into account (including the past of Y). As we had few hypotheses about the directions of connectivity between regions, we choose to use GCM because it does not require a-priori modeling of the connectivity. Given the large amount of regions included in our network, it did not seem feasible to map all possible connections, which would be required for other approaches, such as dynamic causal modeling. We obtained maps of directed influence (dGCM) per session for all regions in the anatomical mask (see Supplementary Materials, Table S1) in each individual participant (N = 20). These dGCM maps are difference maps that have a positive value at voxels where influence *from* the reference region dominates and a negative value at voxels where influence *to* the reference region dominates (Roebroeck et al., 2005). The individual dGCM maps were corrected for multiple comparisons using an FDR of 0.01. The individual dGCM maps were subsequently used in a second level RFX group analysis to calculate differences in directed connectivity between conditions. For the group analysis a RFX ANCOVA, with condition (1PP vs 3PP) as a within-subjects factor, was calculated for each ROI. The resulting F-maps were corrected for multiple comparisons using the cluster level statistical threshold estimator plugin of BrainVoyager QX 2.8 (Goebel et al., 2006) with an initial threshold of p = 0.005 and a cluster size corrected threshold of p < 0.05. In order to counteract a potential downfall of GCM - that directed connections may reflect inherent hemodynamic differences between regions (Deshpande et al., 2010) – we compared effective connectivity *between two conditions* in an *identical set of regions.* If there are inherent hemodynamic differences between regions, which give rise to false connections, these would be present in both conditions. By only taking into account differences in connections between conditions we circumvent the above mentioned problem.

## Results

### Subjective experience of perspective

The results from the questionnaire analysis (Fig. 2) showed that first person perspective training induced higher ratings of Plausibility, Body Ownership and Identification during the perception of threat than the third person perspective training. We assessed the strength of perceived Plausibility, Body Ownership and Identification that participants experienced while viewing the 3D video in the MRI scanner by analyzing the VR questionnaires that were administrated at the end of each session. All questions were scored on a 1 (Not at all) to 7 (Completely) Likert scale. The answer scores were analyzed using an ordinal logistic regression analysis (see Methods section).

Plausibility was assessed with the question “To what extent have you experienced the situation as if it was real?”. The results of the regression analysis (Fig. 2A) showed a significant effect of Session, with higher scores of Plausibility in the 1PP session. The odds of high Plausibility in the 1PP session is 3.337 (95% CI, 1.058 to 10.528) times that of the 3PP session (Wald χ2(1) = 4.226, p = 0.04). Body ownership was assessed with the question: “To what extent did you feel in the female body and lived the situation as if you were the woman?”. The results of the regression analysis (Fig 2B) showed a significant effect of Session, with higher scores of Body Ownership in the 1PP session. The odds of having high Body Ownership in the 1PP session is 3.881 (95% CI, 1.197 to 12.577) times that of the 3PP session (Wald χ2(1) = 5.109, p = 0.02). Identification was assessed with the question: “To what extent did you feel identified with the female body during the experience?”. The results of the regression analysis showed a significant effect of Session, with higher scores of Identification in the 1PP session (Fig. 2C). The odds of having high Identification in the 1PP session is 3.398 (95% CI, 1.056 to 10.935) times that of the 3PP session (Wald χ2(1) = 4.208, p = 0.04).

### Peripersonal space network involved during first person perspective induced threat perception

The results of the fMRI analyses confirmed our first hypothesis: the PPS network was more strongly involved in first than third person perspective primed threat perception. We employed intersubject correlation (ISC; Hasson et al., 2004), a method for investigating brain activity during the perception of natural scenes, to analyze the brain data. In ISC analysis, the shared neural processing of participants is defined by calculating the correlation coefficient between fMRI time series of participants in locations across the brain (see Methods). This makes ISC particularly suitable for naturalistic stimuli, such as 3D video, as it does not require modelling of the stimuli (Hasson et al., 2008; Hasson et al., 2010; Nummenmaa et al., 2012). After calculating an ISC map for each condition, we calculated ISC difference maps, which show the statistical difference between conditions in each voxel (see Methods, N = 20, p[FWER] < 0.05). The analyses were performed within an anatomical mask, which covered most of the cortex (see Supplementary Materials, Figure S1 and Table S1). First person perspective priming induced higher ISC than third person perspective priming in left dorsal and ventral PM, in left IPS, left SMG, bilateral SPL and bilateral PVC during perception of the 3D video (Fig. 3, top). Additionally, after both first (Fig. 3, top) and third person perspective priming (Fig. 3, bottom) differences in ISC were found in different areas of the bilateral PAC, left MT and right PSC.

The results of the ISC difference analyses indicated that during first person perspective primed threat perception regions of the PPS network (PM and IPS) and the SPL and PVC became more synchronized across participants. This suggests that the PPS network is more involved during first than third person perspective primed threat perception. Additionally, the SMG, which is thought to support first person perspective (Maguire et al., 1998; Ruby and Decety, 2001), also showed higher ISC after first person perspective priming. The PSC on the other hand, showed enhanced ISC in both the first and third person perspective primed threat perception.

### Virtual reality training influences effective connectivity in the peripersonal space network

The results from the effective connectivity analyses did not confirm our second hypothesis – that visual and somatosensory regions would send information to PM and IPS during first person perspective primed threat perception. We calculated effective connectivity differences between the two conditions in an identical set of regions using RFX ANCOVA analyses (see Methods, N = 20, p[corrected] < 0.05).

We found that during first person perspective primed threat perception directed connections from PM, IPS, SMG and MT towards SPL were stronger (see Fig. 4 top). This suggests that information was integrated in SPL. These findings are in line with the ISC results, which also emphasized changes in these regions during the first person perspective session. Moreover, we found that bilateral PM showed directed connections to many of the other regions in the network. Additionally, not shown in Fig. 4, we found stronger directed connections from left PAC and left IPS to right ACC and from right ACC to right AMG and left ACC after first person perspective training. Although we found that ISC was reduced in the PPS network during third person perspective primed threat perception, the connectivity results revealed a more complex situation (see Fig. 4 bottom). Contrary to the first person perspective session, we find stronger directed connectivity from IPS to PM, but no integration of information in posterior parietal cortex. Moreover, we found enhanced top-down connectivity from PM, IPS, SMG, SPL and PSC towards visual areas PVC and MT. Additionally, not shown in Fig. 4, we found a directed connection from left SPL to right ACC after third person perspective training.

### Threat processing in nearby space

Finally, we investigated our last hypothesis, that activity in emotion-related structures, such as AMG, is more strongly synchronized across participants during first person perspective experience of nearby threat. We calculated the time course of ISC in the ROIs of three emotion-related structures (bilateral AMG, INS and ACC) using a 10-sample moving average (see Methods). Subsequently, we correlated the time course of the ROI with a predictor that coded for threat distance (low ISC when the aggressor was far away and high ISC when the aggressor was nearby). We found that during the 1PP session the time course of ISC in the amygdala (R = 0.7618, p < 0.0001), ACC (R = 0.7041, p < 0.0001) and insula (R = 0.3519, p < 0.005) correlated with threat distance. In the 3PP session the time course of ISC in the amygdala (R = −0.2137, p =0.07), ACC (R = −0.040, p = 0.74) and insula (R = 0.1623, p = 0.17) did not correlate with threat distance. These results indicate that after first person perspective training threat processing was enhanced when the aggressor was inside the peripersonal space, while this was not the case after third person perspective training.

## Discussion

### Influence of perspective priming on the peripersonal space network

Our first hypothesis was that activation in the PPS network will be stronger when a threat is primed to be in our own space (first person perspective) than in another’s space (third person perspective). The results of the ISC analyses indicated a clear effect of first versus third person perspective priming on the neural activity in the PPS network. After first person perspective priming we found that all regions of the PPS network, including PM, IPS, SMG, SPL and PSC, were more synchronized across participants during PPS intrusion. This was not the case when participants were primed with a third person perspective. We expected that information gathered during threat processing from PVC, PAC, and PSC would converge via the IPS in the PM, as the PM should initiate the defensive responses. We based this hypothesis on the fact that electrical stimulation of F4 and VIP in monkeys (human homologues of superior vPM and ventral IPS) produces movements similar to defensive movements followed by air-puffs (Cooke and Graziano, 2003, 2004). Research in humans also indicated the involvement of PM in threat-related processing (Pichon et al., 2012). Therefore, the ventral PM-IPS network is believed to function to protect the near PPS (Clery et al., 2015). Our findings do indicate the involvement of superior ventral PM and IPS (Fig. 3), but show a convergence of information in SPL rather than vPM. The SPL is an area where many different cognitive functions converge, including attention (Kastner et al., 1999), spatial imagery and perception (Ungerleider and Haxby, 1994; Formisano et al., 2002; de Borst et al., 2012) and the generation and guidance of actions (Culham and Kanwisher, 2001). The superior parietal cortex also underlies the transformation of multisensory input to different coordinate systems, e.g. head, arm, body centered (Andersen et al., 1993), and converting this information into motor commands and whole body actions to targets (Buneo and Andersen, 2006; Whitlock et al., 2008). Together with the PM and IPS the SPL may monitor, predict and evade intrusive actions towards the body (Lloyd et al., 2006; Clery et al., 2015). Our results indicate that this network is more strongly involved during first than third person perspective primed approaching threat perception.

After third person perspective training we observed that brain activity in the PPS network was not consistent across participants. This indicates that after first person perspective training the PPS is aligned with the virtual body and intrusion of this space synchronizes activity in the fronto-parietal network, while after third person perspective training this does not occur. Indeed, our ISC findings suggest that priming with a third person perspective does not activate the defensive PPS network and this aspect of the results did not provide support for a shared PPS. However, the connections between regions of the PPS network did show third person perspective specific modulations. We found an increase of top-down connectivity from the PPS network nodes (PM, IPS, SMG, PSC) towards the primary visual area and MT. This could indicate that increased monitoring of the moving visual stimulus took place after third person perspective priming. Although the threat video that the participants viewed was identical in both conditions the top-down modulation of visual areas we observed here may indicate that the imposed third person perspective altered spatial attention, similar to how task differences can alter attention during identical stimulation (Li et al., 2004).

### Role of the supramarginal gyrus

Our third research question related to the role of the TPJ/SMG in the PPS network. Previous research has shown that the SMG is implied in perspective taking (Falk et al., 2012; Besharati et al., 2016), interoception (Kashkouli Nejad et al., 2015) and self-other distinction (Steinbeis et al., 2015). For example, viewing painful stimulation of a hand evoked activity in bilateral SMG, which was linked to simulated pain to the own body (Costantini et al., 2008). The SMG has also been mentioned as part of the multisensory representation of PPS (Lloyd et al., 2003; Makin et al., 2007; Brozzoli et al., 2012a; Grivaz et al., 2017). The results of our ISC analyses indicated that the SMG is more strongly involved in first than third person perspective primed threat perception. Moreover, during first person perspective primed threat perception the SMG was an integrated part of the PPS network: it received information from PM and sent information to SPL. These results suggest that the SMG plays a role in relating body threatening information to the self. In the 3PP session brain activity in the SMG was not strongly synchronized across participants. It did, however, show top-down connections to the visual areas.

### Peripersonal space and threat perception

Our final research question focused on synchronized brain activity in emotion-processing regions during first person perspective experience of nearby threat. Defensive responses are especially enhanced when the threat is near, as shown by a series of experiments with virtual characters (Ahs et al., 2015) and threatening stimuli (Mobbs et al., 2009; Mobbs et al., 2010; de Haan et al., 2016; Wabnegger et al., 2016). A recent meta-analysis showed that the amygdala is more activated when moving from threat anticipation to threat confrontation (Klumpers et al., 2017). Our results support these findings. We found that brain activity is more synchronized across participants in AMG, INS and ACC when threat is near, but only when the threat is perceived as directed to one-self (1PP condition). We found no evidence for enhanced ISC for nearby threat in emotion processing regions when the threat was perceived as directed to another person. Together with the behavioral findings this indicates that the virtual reality training from first person perspective was effective in eliciting identification with the virtual victim and enhancing affective responses to nearby threat (in line with Galvan Debarba et al., 2017).

In addition we found modulations of connectivity to the emotion processing regions during the perception of the threat scenario both after first and third person perspective training. The connectivity analyses revealed that in the first person perspective session PAC and IPS sent information to right ACC and from the right ACC information was forwarded to right AMG and left ACC. The ACC has a general role in decision making and social and emotional processing (see Lavin et al., 2013) and has bilateral connections to the premotor and auditory cortex (Jacobson and Trojanowski, 1977; Muller-Preuss et al., 1980; Vogt and Pandya, 1987; Barbas, 1988; Petrides and Pandya, 1988; Barbas et al., 1999). Moreover, the ACC is linked to vocalizations (Muller-Preuss and Ploog, 1981; Dolan et al., 1995; Frith and Dolan, 1996) and auditory processing of speech in humans (Paus et al., 1993; Frith et al., 1995) and is connected to the limbic system, including the AMG. This link suggests a role of ACC in the appraisal and regulating of emotions (Etkin et al., 2011) and in encoding emotional significance of auditory stimuli.

### Future directions

In this study we found that virtual reality priming is particularly suitable for fMRI experiments. Our study exemplifies how priming of body ownership in VR virtual reality outside of the MRI environment, in combination with a 3D video during fMRI measurements, can greatly contribute to the possibilities of naturalistic methodologies in social and affective neuroimaging experiments. The scarcity of human neuroscience studies using dynamic threatening stimuli contrasts with the relevance of personal space intrusion in the perception of threat, both for the victims (Lloyd et al., 2006) and for the aggressors (Schienle et al., 2016). As static threatening stimuli may fail to evoke realistic responses, naturalistic immersive 3D motion stimuli appear better suited to modulate the distance of threatening stimuli within a naturalistic environment that the participants may perceive as real. Our study shows that first person perspective visuo-motor training in virtual reality can be utilized to prime subsequent experiences such that behavioral and neuronal responses are aligned with the virtual victim. Combining immersive virtual reality training with neuroimaging methods could provide a basis to change perspective during therapeutic treatment (Seinfeld et al., 2018).

## Acknowledgements

We would like to thank Guillermo Iruretagoyena, Xavi Navarro, Sofía Seinfeld, and Chris Wiggins for their help with the virtual reality stimuli and set-up and Giancarlo Valente and Jukka-Pekka Kauppi for their suggestions on the ISC analyses.

This work was supported by FP7/2007-2013, ERC grant agreement number 295673.

